# Identification of lactate as a potential inhibitory factor within epithelial cell supernatant and its effects on type 2 T cell populations

**DOI:** 10.1101/2020.03.26.008292

**Authors:** Francis Hughes, Ian D Pavord, Timothy J Powell

**Author notes:** Corresponding authors Dr Timothy Powell. Prof Ian Pavord.

## Abstract

Epithelial cells have been shown previously to inhibit release of type 2 cytokines from T cells and this process could be important in a type 2-mediated diseases such as asthma. Factors secreted by epithelial cells could regulate inflammatory responses and release an ‘all is well’ signal such that the immune system is controlled. Here we show that lung epithelial cells are able to inhibit IL-2 driven type 2 cytokine release by PBMC cells across a transwell indicating the involvement of soluble mediators. We then fractionate the mediators within the conditioned media using methanol chloroform separation and size fractionation. Inhibitory activity was demonstrated within the polar fraction of supernatant separated by methanol chloroform extraction. We then found that lactate, a component within the polar fraction was able to mediate inhibition of the type 2 cytokines. The inhibitory activity of lactate deserves further study and could play a role in the inhibition of T cell derived cytokines in vivo.

## Introduction Results and discussion

*To the editor*,

Type 2 immune responses characterised by high levels of type 2 cytokine (IL-4, IL-5 and IL-13) secreting T cells, eosinophils, basophils and cytokines such as IL-4, IL-5 and IL-13 are key drivers of immunopathology in the airways of the dominant phenotypes of asthma [1]. Inhibition of Th2 cells by epithelial cells has been shown *in vivo* and *in vitro* [2, 3] and may be important in regulating the pathogenesis of asthma [4, 5]. We have characterised an experimental system (see materials and methods online supplement) using peripheral blood mononuclear cells (PBMC), taken from healthy cell bank donors, cultured with IL-2 in the presence and absence of epithelial cells. We found that the presence of A549 alveolar, BEAS2B or 16HBE bronchial epithelial cells led to reduction in IL-2-driven IL-13 and IL-5 release across a transwell and this inhibition is epithelial cell number dependent (Figure 1A). Similar data were seen with healthy human bronchial epithelial cells (HHBEC) cultured with PBMC incubated with IL-2 (Supp Figure 1A). IL-2 driven release of other cytokines such as TNFα were also inhibited by epithelial cells indicating that this inhibitory process may not be exclusive to type 2 cells (data not shown). PBMC were then cultured with IL-2 and epithelial cell conditioned media (CM) which reduced the amount of IL-13 (Figure 1B) or IL-5 (Supp. Figure 1B) secreted by the PBMC in a CM dose dependent fashion. Then we compared the ability of CM derived from a high number (5 × 10^5^) or a lower number (5 × 10^3^) of epithelial cells in a 24 well plate well. Conditioned media from 5 × 10^5^ cells showed greater inhibitory effect in a dose dependent fashion than CM from 5 × 10^3^ cells (Figure 1C, for absolute values see Supp Fig 2A). We then separated the supernatant into size specific fractions and found that fractions less than 3kDa were similar in their inhibitory activity to those above 3kDa (Supp Fig 2B). Further separation of the supernatant using methanol / chloroform extractions showed that the protein fractions were less inhibitory than the polar and organic fractions, with the polar fraction being the most-inhibitory (Figure 2A). Combined size fractionation and methanol / chloroform extraction resulted in the polar fractions being most inhibitory whilst size fractionation seemed less critical (Figure 2B). The supernatant from the epithelial cell / PBMC co-cultures was generally more acidic than that from PBMC alone which indicated more metabolic activity. Lactate was a possible mediator because it is located within the polar fraction of the supernatant and has been shown to be associated with higher numbers of cells in culture [6]. We then cultured PBMC in the presence of IL-2 with increasing levels of sodium lactate and lactic acid and found that the release of IL-13 and IL-5 was reduced in the presence of lactate and lactic acid (Figure 2C). The lactic acid clearly acidified the media as indicated by pH indicator colour change, however sodium lactate inhibited the type 2 cytokine release without an associated acidification of the media. Other investigators had shown that epithelial cells could inhibit mast cells via a pertussis toxin sensitive pathway [7]. Addition of pertussis toxin showed that this inhibitor reduced the ability of lactate to prevent IL-13 or IL-5 release from the PBMC via a lactate concentration-dependent mechanism (Figure 2D). Thus lactate may mediate inhibition of type 2 cytokines through a G-protein sensitive pathway.

**Figure 1:**
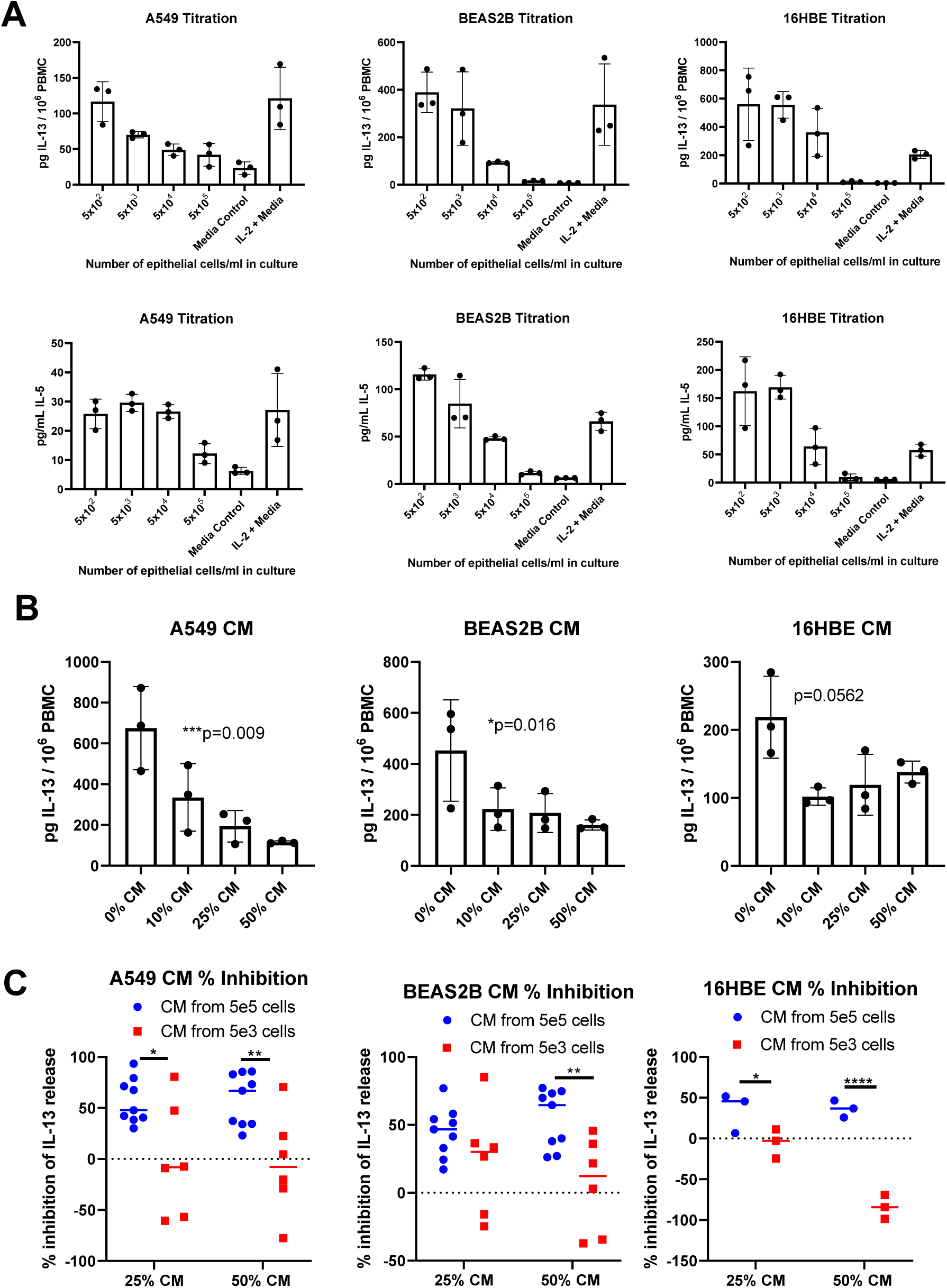
IL-2 driven secretion of IL-13 and IL-5 from PBMC is regulated by epithelial cells and conditioned media. A) Titrated numbers of epithelial cells were adhered to 24 well plate wells overnight and the next day 10^6^ PBMC in transwells plus IL-2 (200 U/mL) were added on top. Five days later supernatants were collected and cytokine content measured by ELISA. Upper row IL-13, lower row IL-5. Data shown are means +/- SD. Data are from one representative experiment repeated at least twice. B) Conditioned media from epithelial cells were added at titrated amounts to 96 well plates then 2 × 10^5^ PBMC added plus IL-2 and five days later content of IL-13 measured. Data shown are means +/- SD with statistical comparison of linear trend. Data are from one representative experiment repeated three to six times. C) Similar to B, CM experiments were performed and IL-13 measured and then percentage inhibition calculated. Statistical comparisons are two-way ANOVA followed by Sidak’s multiple comparison test. Data shown are from six experiments (A549, BEAS2B) or three experiments (16HBE) and each data point represents a mean of three individual wells.

**Figure 2:**
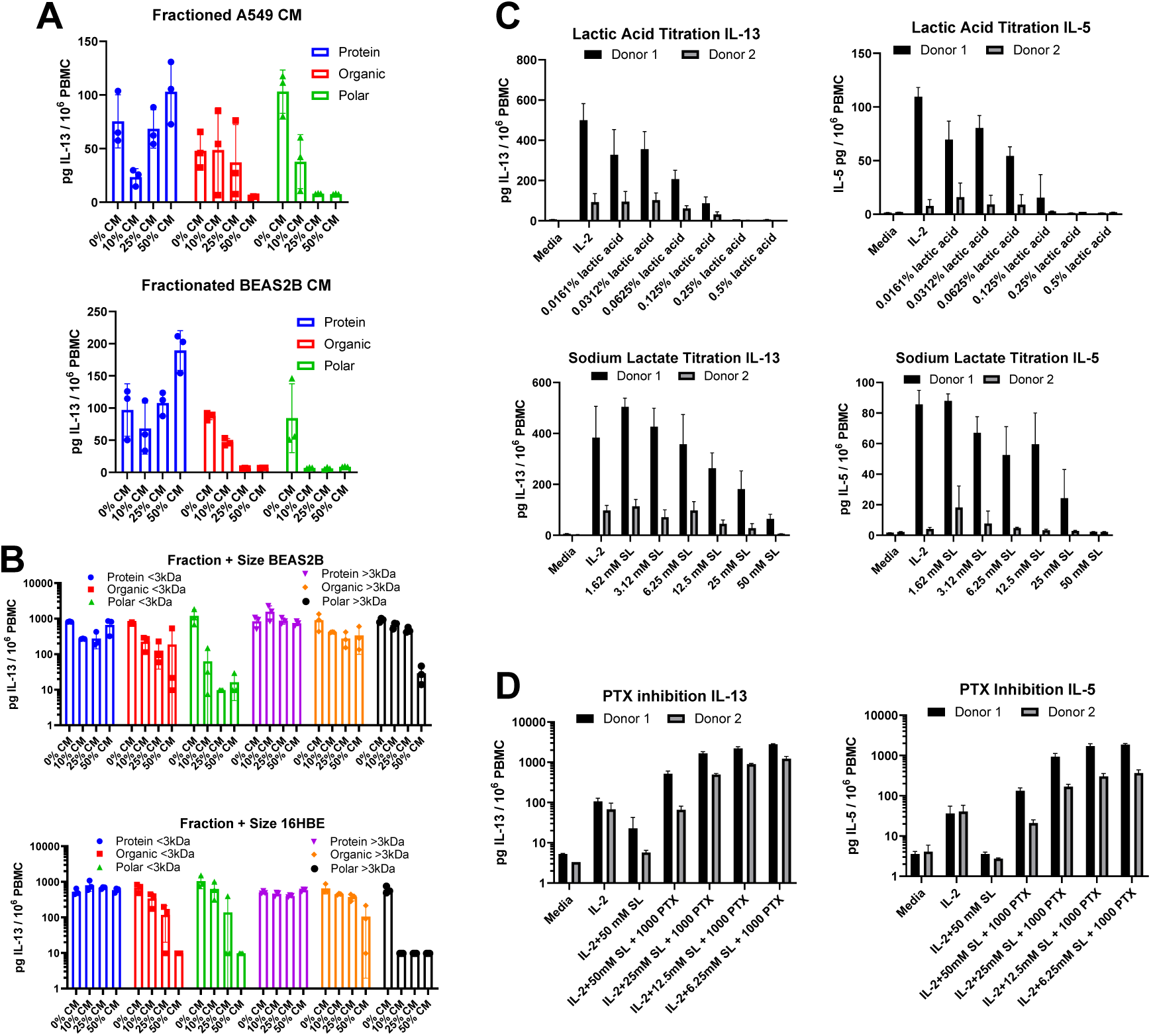
Separation of conditioned media indicates inhibition is mediated by a polar molecule lactate which in turn can be inhibited by pertussis toxin. A) Supernatants were fractionated into protein, organic and polar fractions using methanol / chloroform separation and then cultured in titrated doses with PBMC and IL-2. Five days later supernatants were harvested and IL-13 content measured. Values shown are means +/- SD of three wells. B) Supernatants were separated into size fractions followed by methanol/chloroform extractions and resultant fractions cultured with PBMC and cytokine content measured after five days. Values shown are means +/- SD of three wells. C) PBMC were cultured with titrated doses of lactic acid and sodium lactate. IL-5 and IL-13 concentrations were measured 5 days later. Values shown are means +/- SD using two different PBMC donors. D) PBMC were cultured with lactate in the presence or absence of pertussis toxin at the concentration shown. IL-13 and IL-5 were measured by ELISA five days later. Means +/- SD are shown with two different donor compared to C). Data are from one representative experiment repeated at least twice.

The release of inhibitory factors by epithelial cells could be an important immunoregulatory mechanism in asthma, particularly in the context of epithelial cell barrier damage and dysfunction in asthma. Although epithelial dysfunction results in stimulatory factor production it is also possible that inhibitory factors, including this novel lactate dependent mechanism we describe, are reduced [2, 3, 7]. The epithelial cell has been shown to inhibit T cells via a soluble pathway and the nature of these mediators has yet to be established [8]. Here we show that the polar fraction of the epithelial cell supernatant is most inhibitory and a potential factor from that fraction is lactate. It is unlikely that lactate is the only inhibitory substance within the epithelial cell supernatants and it is possible that different epithelial cell lines secrete different moieties. Lactate has been shown by others to inhibit T cell motility and this, again, is in addition to the simple acidification by lactic acid [9].

However it is very compelling that lactate and general metabolism is important for the inhibition of these potentially pathogenic type 2 T cells and this may open up novel therapeutic pathways for further investigation.

## Acknowledgements

We would like to thank Dr Timothy Hinks, Mary Brown and Sophie Morgan and other members of the respiratory medicine unit for experimental input, helpful discussion and comments on the manuscript. We also thank Prof Benedikt Kessler from the Target Discovery Institute for helpful discussion and experimental advice.

## Supplementary Materials and Methods

### Blood samples

Blood cones were from consented healthy donors and were supplied using NHS blood and transfusion (NHSBT) ethics permission T298. Blood cones were obtained from the NHS blood and transfusion service and then PBMC separated on density gradients using lymphoprep (Stemcell). All donors provided written informed consent.

### Reagents and materials

The cytokine IL-2 was purchased from Peprotech and sodium lactate was from Sigma. Lactic acid (Scientific Laboratory Supplies), methanol and chloroform were Analar grade from Sigma. Bordetella pertussis toxin (PTX) full length protein was purchased from Abcam and diluted in X VIVO media and used at a final concentration of 1000 ng/mL.

### Cell Lines and epithelial cell supernatant

A549 were obtained from Prof Luzheng Xue (Respiratory Medicine Unit) being originally purchased from the ECACC and were cultured in D10 media, DMEM media (Sigma) with 10% v/v foetal calf serum (FCS) (Sigma), 2mM glutamine and 100 U/ml penicillin and 100 ug/mL streptomycin (Gibco). BEAS2B and 16HBE cells were a gift from Prof Ling-Pei Ho (University of Oxford) and were cultured in F10 media, Hams F12K media (Gibco) with 10% v/v FCS, glutamine and penicillin / streptomycin as above. Supernatant was generated from 5 × 10^5^ or 5 × 10^3^ epithelial cells per well, plated into 24 well plates overnight. Then media was replaced with serum free X-VIVO 15 media (Lonza) and cells were cultured for 48 hours and supernatant clarified by centrifugation and filtered through 0.2um filters.

### Human Bronchial epithelial cells (HBEC)

HBEC were obtained from human bronchoscopy samples and cultured in complete HBEC media (Lonza) and put into 24 well plates and used when 90% confluent.

### IL-2 driven cytokine secretion assays

PBMC were cultured in X-VIVO 15 media (Lonza), in the presence of IL-2 (200 U/mL) for five days in an upper transwell with either media or confluent epithelial cells in a 24 well plate below the transwell. After the culture period supernatants were harvested and IL-13 (Thermo eBioscience Ready set go) or IL-5 (Mabtech ELISA development kit) concentration measured by ELISA. For cell culture supernatant conditioned media (CM) inhibition experiments, titrated amounts 10%, 25% or 50% v/v of CM were added to 96 well plate bottom plates and a fixed number 2 × 10^5^ PBMC with 200 U/mL IL-2 as appropriate. Five days later supernatants were collected and cytokines measured as above. Sodium lactate was added at titrated doses between 1.62 and 50 mM and lactic acid added at % v/v between 0.5% and 0.0161% throughout the course of the assay. PTX was tested by titration (not shown) and added at a final concentration of 1000 ng/mL for the course of the assay.

### Separation of supernatants into fractions

Epithelial cell conditioned media (CM) was separated using size exclusion centrifugation devices (Millipore) such that supernatants were added to the top of the tubes above the membrane and centrifuged at 5000g for 1h at 4 °C. Then <3KDa supernatant was collected directly from the lower part of the tube. Supernatant >3KDa that remained above the membrane was diluted 1:10 with X-VIVO so that the original volume was regained. These supernatants were then cultured in titrated doses 10%, 25% or 50% v/v with PBMC as above. For protein, organic and polar separation, 200 µL supernatant was mixed with 600 µL methanol and 150 µL chloroform, vortexed, then 450 µL sterile H_2_O added and vortexed again. The mixture was then centrifuged at 13000g for 1 minute. The upper aqueous phase was pipetted off and kept as a polar fraction. Then 450 µL methanol was added to the protein/organic phases and vortexed. After centrifugation for 2 minutes at 13000g supernatant was removed to form the organic phase and the pellet was kept as the protein phase after resuspension in 200 µL X-VIVO. The polar and organic phases were centrifuged in a concentrator plus (Eppendorf) to a small volume and then resuspended into 200 µL X-VIVO for polar phase and X-VIVO with 0.5% DMSO for the organic phase. For two stage, size and phase separation, CM was separated into size fractions followed by methanol/chloroform extraction.

## Figure Legends

**Supplementary Figure 1:**
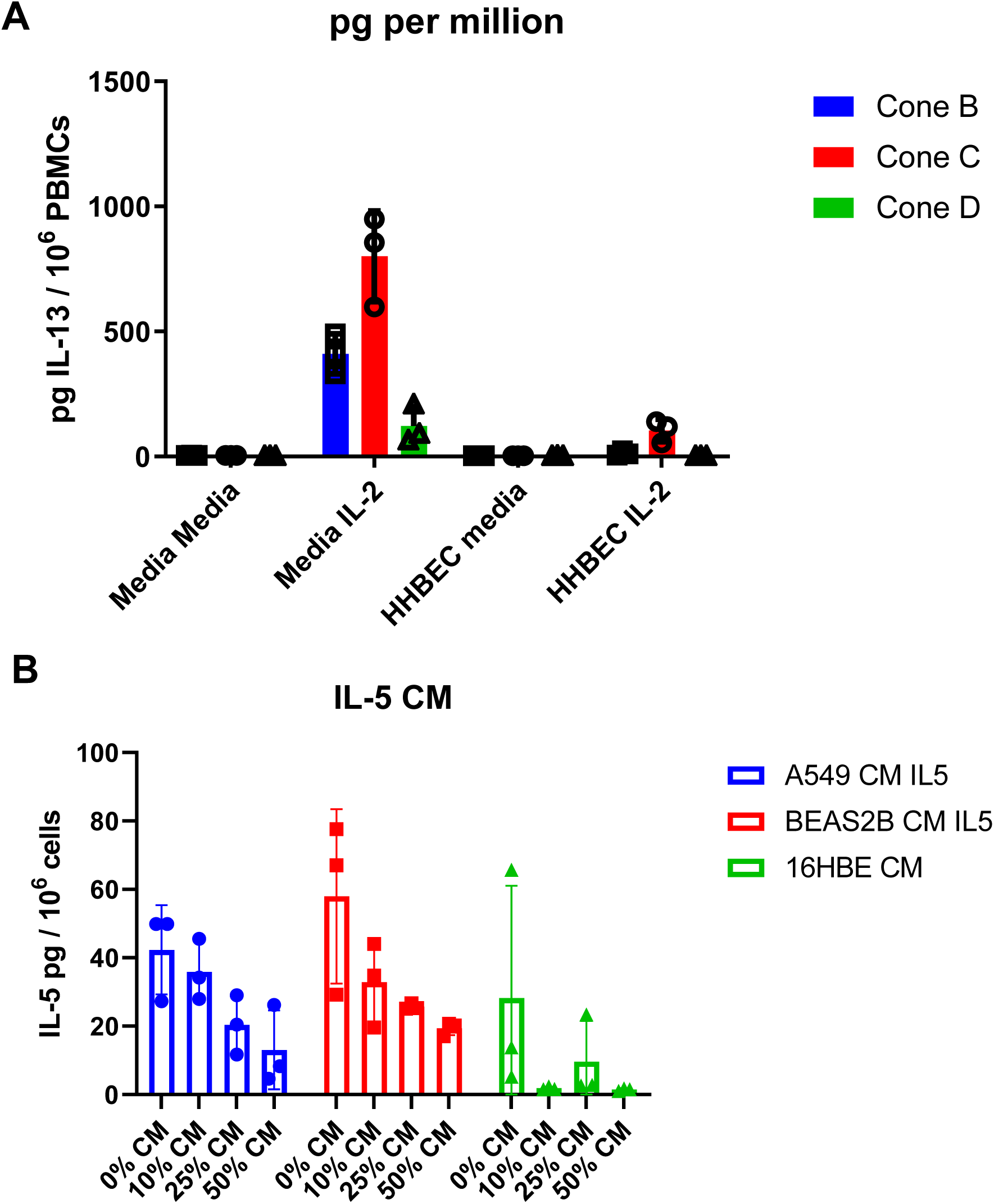
IL-13 release driven by IL-2 is inhibited in the presence of HBEC cells and epithelial cell CM inhibits release of IL-5 from IL-2 stimulated PBMC. A) Human Bronchial Epithelial cells (HBEC) were cultured in 24 well plates, then PBMC (10^6^) with IL-2 (200 U/mL) were added in transwells on top. Media controls were 24 well plate wells containing media alone. After 5 days supernatants were collected and IL-13 measured. Values are means +/- SD from three separate wells and three different PBMC donors. B) PBMC were cultured with titrated doses of epithelial cell CM and after five days culture media collected and IL-5 concentration determined. Values are means +/- SD. Data are from one representative experiment repeated at least twice.

**Supplementary Figure 2:**
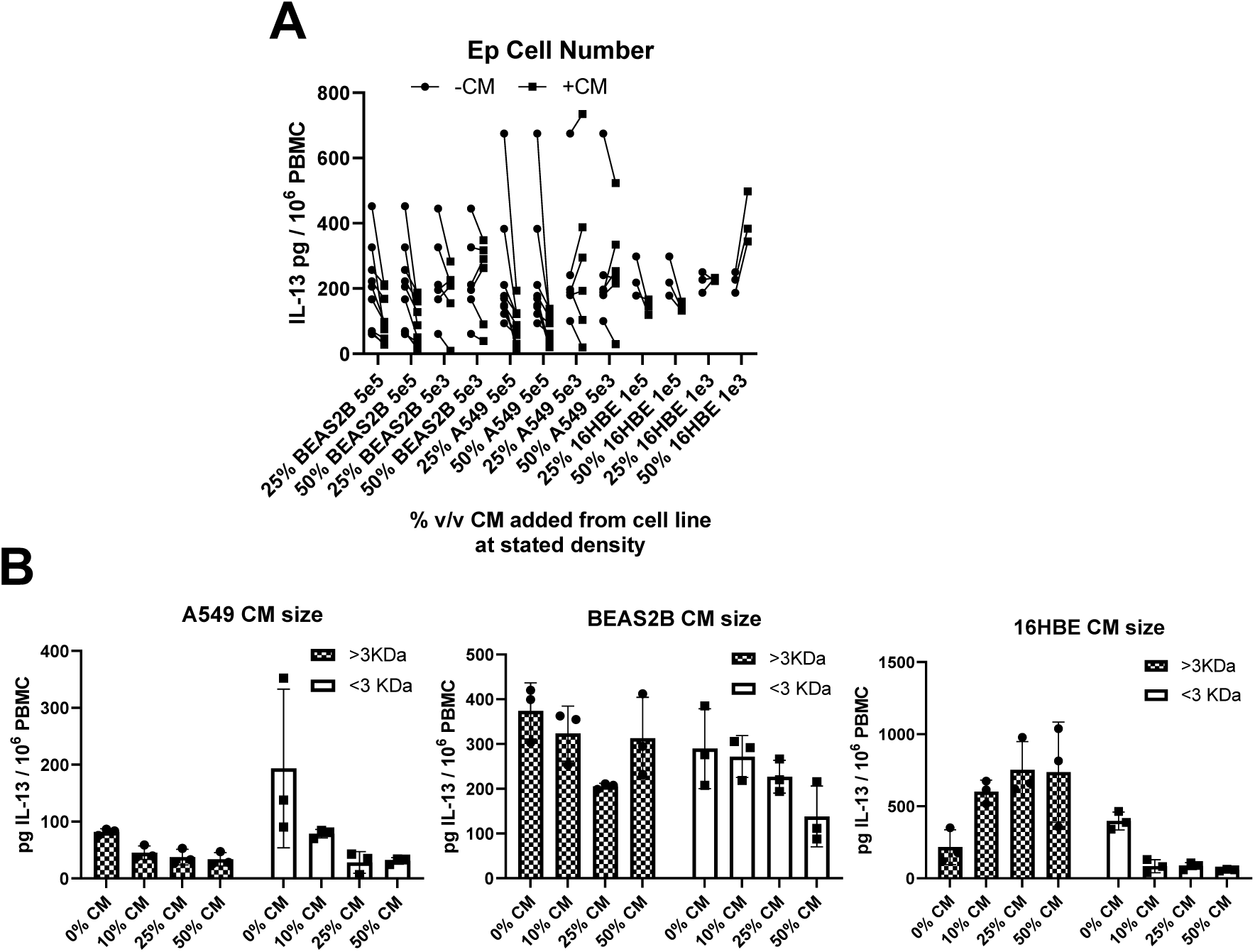
Epithelial cell CM inhibits release of IL-13 but size separation results in similar levels of inhibition. A) IL-13 release was measured in wells in the presence or absence of 25% or 50% v/v CM from A549, BEAS2B or 16HBE cells. –CM represents IL-2 only plus PBMC and +CM means IL-2 plus PBMC +25% or 50% v/v CM from epithelial cells. These values are shown as percentage inhibition in figure 1C. B) PBMC plus IL-2 were cultured with titrated volumes of epithelial cell CM separated previously by size using size exclusion centrifugal filters. IL-13 release was measured five days later. Values shown are means +/- SD.

